# Retinal Vessels Detection Using Convolutional Neural Networks in Fundus Images

**DOI:** 10.1101/737668

**Authors:** Md. Mohaimenul Islam, Tahmina Nasrin Poly, Yu-Chuan (Jack) Li

## Abstract

Computer-aided detection (CAD) system is a realistic option for physicians to screen fundus images. Automated segmentation of retinal vessel is in fundus important step to identify the retinal disease region. However, identification of the retinal disease region accurately is still challenging due to the varied distribution of blood vessel on noisy and low contrast fundus images. Healthcare system has been changing significantly with the emergence of machine learning (ML), deep learning (DL) and artificial intelligence (AI) in recent year. Retinal vessel detection is one such area of application of deep learning, for improving the accuracy of detection and segmentation and the quality of patient care. Recently, the convolutional neural networks (CNN) have been applied to the detection of the retinal vessel from fundus images and have demonstrated promising results. The range of accuracy of the CNN model was 0.91-0.95 and the area under the receiver operating curve was 0.09-0.98. Therefore, CNN may play a crucial role in determining the therapeutic methods and detecting the retinal vessel accurately in an individual manner. In this survey, we described the use of CNN in fundus imaging, especially focused on CNN technique, clinical application for retinal vessel detection and future prospective.

## Introduction

Recently, deep learning algorithms have been extensively used in almost every area in healthcare because of their successful outcomes in image recognition, natural language processing, speech recognition, bioinformatics, computer vision, cancer, diabetic retinopathy^1, 2^. The identification of interaction between drugs, development of new drug molecule, diagnosis of cancer and other diseases, automated treatment and dose recommendation is a very complicated process, but deep learning algorithms have created new hope that they could aid in helping physicians make the right considerations in diagnosis and treatment by providing as an immense of scientific knowledge^3, 4^. It is also contributing to biomedicine including computational drug discovery. By using the tremendous deluge of healthcare data, deep learning algorithms are actually helping physician to make real-time clinical decision making for their patients.

Retinal vessel detection is an appropriate application for precision medicine and artificial intelligence techniques because of monitoring retinal vessel in fundus images may assist physicians to correctly diagnosis and treatment of ophthalmologic diseases ^5^. As manual detection of the retinal vessel is challenging and time-consuming; therefore, there is an unmet and urgent need to develop an automatic detection of retinal vessel for cost-effective screening and treatment. The use of the CNN algorithms in the retinal vessel detection has generated much interest over the few years. This review aims to evaluate the effectiveness of the CNN model for detection of retinal vessels.

### Deep learning

Originally developed as mathematical theories of the information-processing activity of biological nerve cells, the structural elements used to describe an ANN are conceptually analogous to those used in prediction model/ recommendation model, despite it belonging to a class of statistical procedures ^6^.The basic element of the model is given below:

### Perceptron

It is a simple algorithm that takes an input vector x of m values (*x*_1_, *x*_2,_ …. . *x*_*n*_) which is often recognized as input features or simply features. It then multiplies them by some factors called “weight”, represented by *w*_1,_*w*_2_, …. *w*_*n*_, gives an outputs either 1 (yes) or 0 (no).

Mathematically expression of this function is given below:

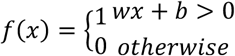

Here, *w* is a vector of weights, *wx* is the dot product of 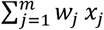, and *b* is a bias. A weight is a connection between neuron that carries a value. However, the higher the value, the larger the weight is. However, *wx* + *b* always defines a boundary hyperplane that changes position according to the values assigned to *w* and *b*. If *x* lies above the straight line, then the answer is positive, otherwise it is negative. Moreover,

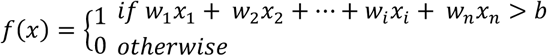

The perceptron often takes some *linear combination of input values or features* compare it to a *threshold value b* and *return* 1 *if the threshold exceeded and zero if not*.

### Multilayer perceptron (MLP)

An MLP (or Artificial Neural Network) is a deep, artificial neural network takes more than one perceptron. An MLP with three-layers, first layer consider as a *input layer*, and last consider as a *output layer* and middle layer consider as a *hidden layer. Input layer* receive input value such as *x*_1_, *x*_2_, …. *x*_*n*_ and take the output (*y*_*n*_) from the *output layer*. However, *number of hidden layer* can be added according to need. An example of a MLP is given below:

It always applies to supervised learning problems in which a set of input and output variables are trained and learn the pattern between those in- and outputs. However, training process involves with adjusting the input parameters or the weights and biases. Because, all the edge weights are randomly assigned initially. The adjustment process need to be repeated until the output error is below or equal to a predetermined threshold. It is, however, important to minimize the errors during the training process. To minimize the errors during the training process, *backpropagation* is used to adjust those weight and biases to make less errors. The error can be measured in a variety of ways such as *root mean squared error* (*RMSE*), *mean bias error* (*MBE*).

In the classification outcome, activation function like *softmax function* is used in the *output layer* to make sure that the outputs are probabilities and they take value up to 1. An arbitrary real-value score is always taken by the *Softmax activation* function and it then convert it to a vector values between zero and one. It is like-

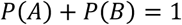

### Activation function

Activation function is an integral part of neural network that is often known non-linearity, describe the input and output relations in a non-linear way. However, non-linearity element allows for higher flexibility and make a complex function during the whole model learning process. It helps to speed up the whole learning process. Several activation functions such as sigmoid, tanh, ReLU are commonly use in practice.

Above activation function *σ* with the (*x*_1_, *x*_2_, … *x*_*n*_) input variables (*w*_1_, *w*_2_, *w*_*n*_) weight vector, *b* bias, and *Σ* summation is given in the following diagram.

#### a) Sigmoid function

It takes a real-value input and convert it to range between 0 and 1. The sigmoid function is defined as follows:

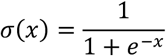

Here it is clear that it will convent output between 0 and 1 when the input varies in (−∞, ∞). A neuron can use the sigmoid for computing the nonlinear function *σ*(*y* = *wx* + *b*). If *y* = *wx* + *b* is very large and positive, then *e*^−*y*^ → 0, so *σ*(*y*) → 1, while *y* = *wx* + *b* is very large and negative *e*^−*y*^ → ∞, so *σ*(*y*) → 0.

b) **Tanh**: It takes a real-value input and convert it to the range of −1 and 1. The tanh function is defined as follows:

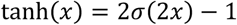

c) **ReLU**: It is called Rectified Linear Unit and takes a real input variable and thresholds it at zero (replace native values with zero). The ReLU function is defined as below:

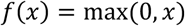

### Loss function

It represents how well specific algorithm model works. *Loss function* would be large if prediction results are far from actual results. Optimizer function is used to reduce the error in prediction. However, *loss functions* can be classified into two major categories such as *regression losses* and *classification losses*. It can be mathematically expressed by-

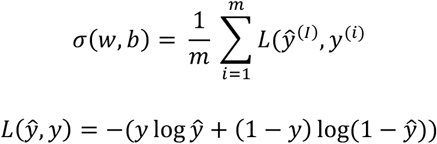

### Optimization algorithms

It is generally use to minimize errors, make slightly better and faster results by updating the input parameters such as weight and bias values. *Gradient Descent* is most widely used optimization algorithms that help to tell us whether the function is decreasing and increasing at a particular point. The following figure represents how *cost function C*(*w*) is one single variable *w*.

The *cost function C* is the initial value and desired point ism *minimum C*_*min*_. However, the starting weight is *w*_0_ and each step represents by *r* and the gradient is the direction of maximum increase. The direction of the value can be expressed mathematically to the partial derivative of 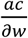 evaluate a point *w*, reach at step *r*. However, the opposite direction can be expressed by 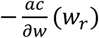. Most commonly used optimizer are *Momentum, Adagrad, AdaDelta, Adam*.

### Convolutional Neural Network

The convolutional neural networks (CNNs) learn to perform their tasks through repetition and self-correction. A CNN algorithm consists of three layers such as convolutional layer, pooling layer and fully connected layer.

### Convolutional layer

It is the main building block of the CNN. It carries the main portion of the network’s computational load. A convolutional layer consist of many *kernels* that consider as a set of learnable parameters. However, it is spatially smaller than an image but is more in-dept. Neuron in the first convolutional layer are not always linked to each pixel in the input image but only to pixels in their receptive fields. However, each neuron in the second convolutional layer usually connect only the neuron located within a small rectangle in the first layer. In this process the network concentrate on low-level features in the first hidden layer, afterwards assemble all into higher-level features in the next hidden layer and so forth so on. The is the most common hierarchical structure that’s why CNNs perform well for image recognition. The formula is given below:

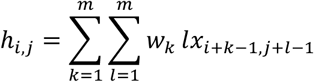

Here, m= kernel width and height, h = convolutional output, x = input and w = convolutional kernel.

### Pooling layer

A pooling layer typically works on every input channel independently, so the output depth is the same as the input depth. It replaces the output of the network at certain locations by deriving a summary statistics of the nearby outputs. Pooling layer helps in reducing the spatial size of the representation that usually help to reduce the computation load and the memory usage. A pooling neuron has no weights; is aggregate the inputs using an aggregation function like the max and mean. In mathematically,

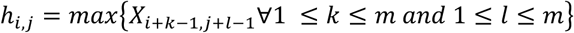

### Fully connected layers

It usually combine features learnt by different convolutional kernel. **Figure 6** shows how it works for detection of retinal vessels.

**figure 1:**
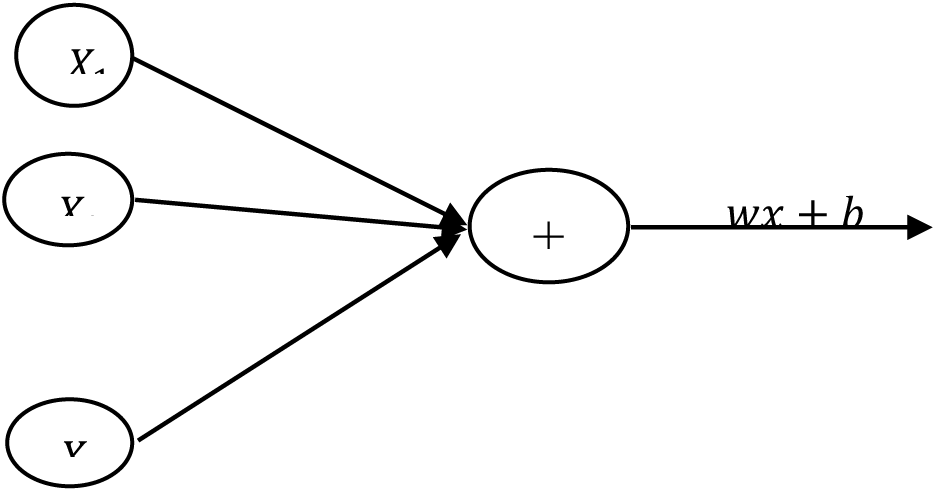
perceptron

**figure 2:**
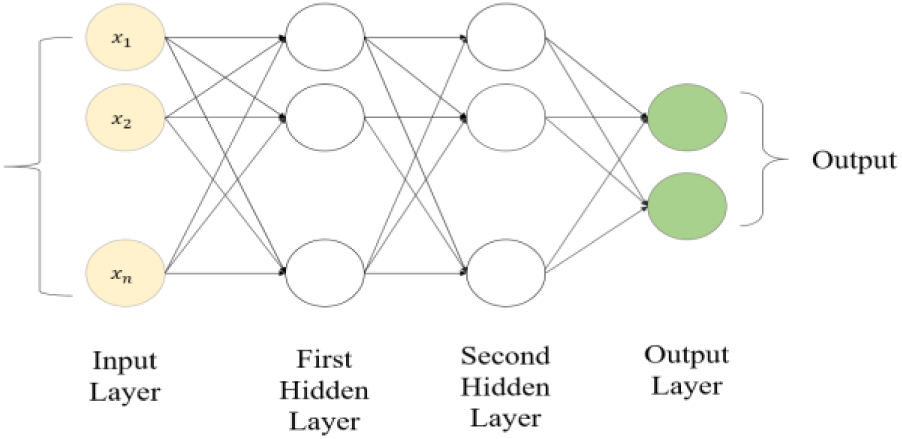
Basic structure of ANN model

**figure 3:**
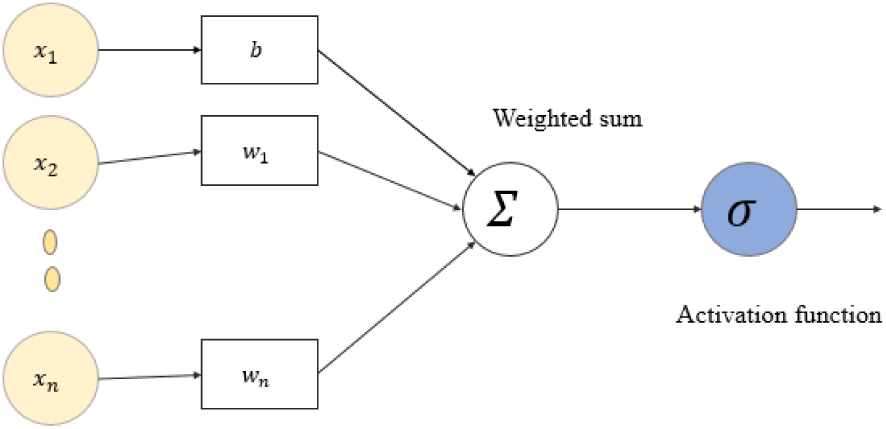
Basic structure of activation function

**figure 4:**
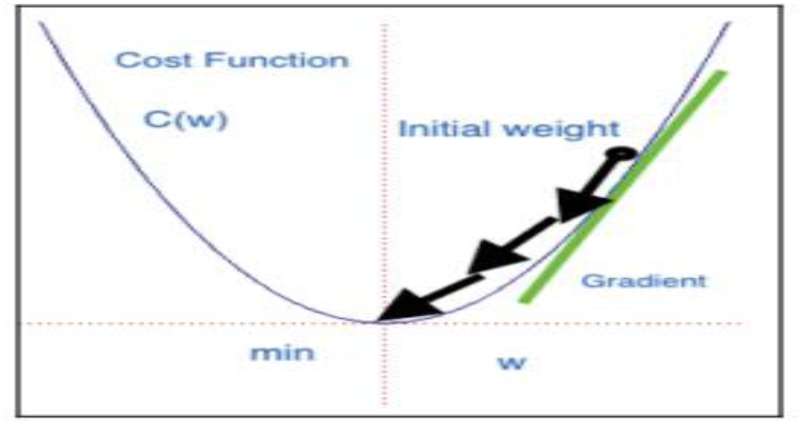
Gradient descent

**figure 5:**
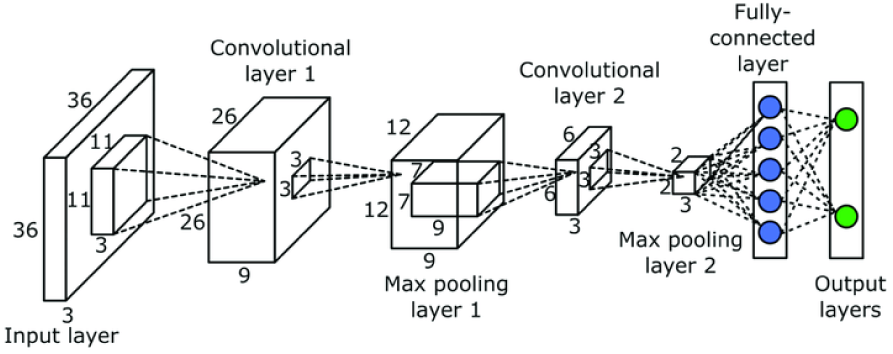
Architecture of CNN

**figure 6:**
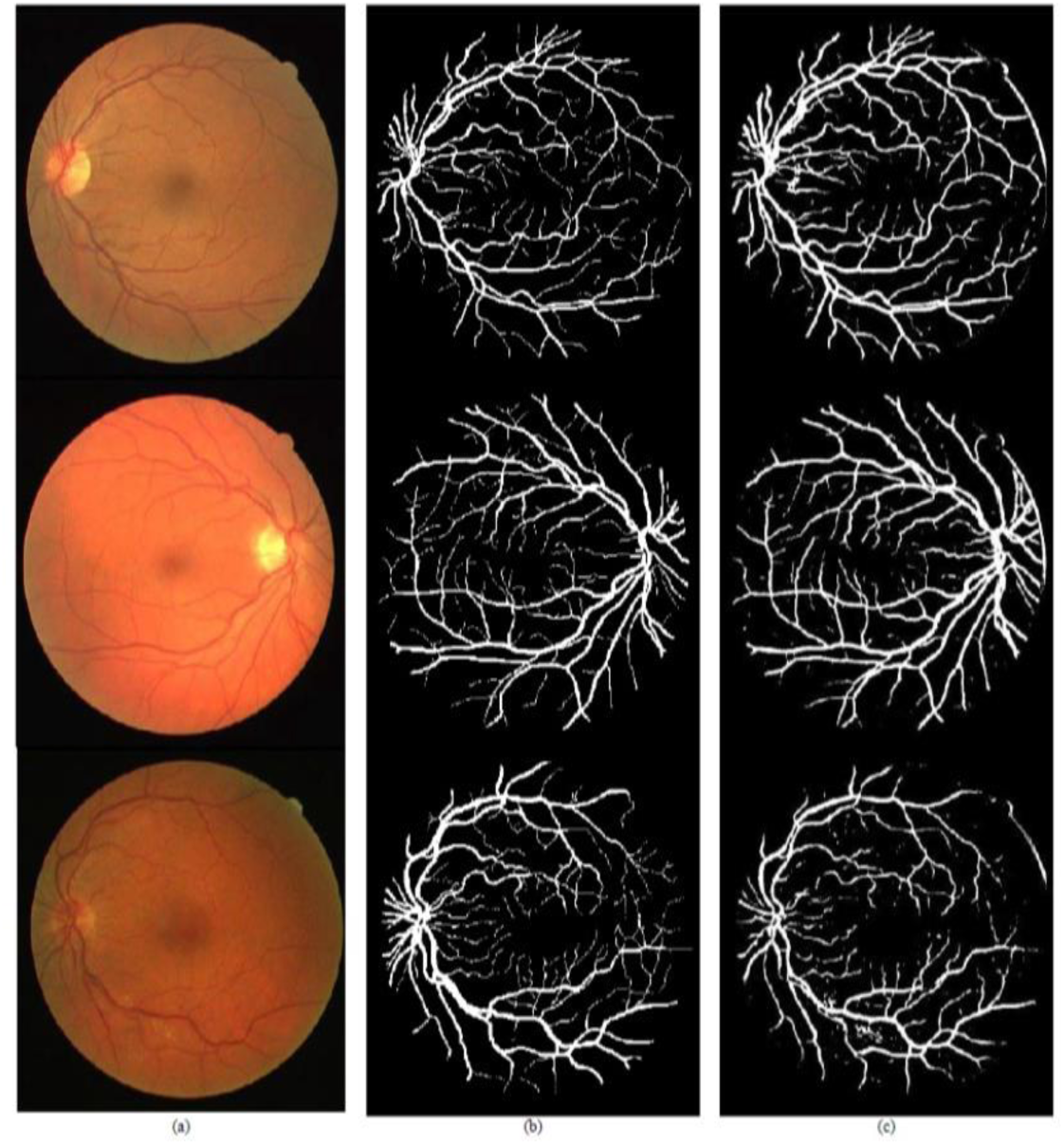
Detection of retinal vessel using CNN a) Original images b) corresponding ground truth and c) detection results.

### Clinical applications

The convolutional neural network could remarkably change the way of retinal disease diagnosis and management in the near future. Automated diagnosis of retinal vessel might be popular in an era where physicians, patients, healthcare policy makers would want to get cost-effective and efficient care. Deep learning based automated detection of retinal vessel would screen patients without requiring any experts to interpret retinal images. Therefore, this could be used by any kinds of physicians who are not normally ophthalmologists. The convolutional neural network algorithms have been applied to detect retinal vessel. Included studies used the DRIVE, STARE, and CHASE DB1 dataset to develop and evaluate their performance. Studies were published between 2015 and 2018. The range of accuracy was 0.91 to 0.95 and the rage of area under receiver operating curve was 0.95 to 0.98. **Table 1** shows the performance of CNN model for retinal vessel detection.

**Table 1:** The performance of CNN model of detection retinal vessel.

| Author | Year | Model | Dataset | SN/SP | Accuracy | AUROC |
| --- | --- | --- | --- | --- | --- | --- |
| Guo<sup>7</sup> | 2018 | MDCNN | DRIVE | 0.98/0.70 | 0.96 | 0.97 |
|  |  |  | STARE | 0.98/0.56 | 0.95 | 0.95 |
| Oliveira<sup>8</sup> | 2018 | CNN | DRIVE | 0.80/0.98 | 0.95 | 0.98 |
|  |  |  | STARE | 0.83/0.98 | 0.96 | 0.99 |
|  |  |  | CHASE DB1 | 0.77/0.98 | 0.96 | 0.98 |
| Dasgupta<sup>9</sup> | 2017 | CNN | DRIVE | - | 0.95 | 0.97 |
| Şengür<sup>10</sup> | 2017 | CNN | DRIVE | - | 0.91 | 0.96 |
| Li<sup>11</sup> | 2016 | CNN | DRIVE | 0.7/0.98 | 0.95 | 0.97 |
|  |  |  | STARE | 0.77/0.98 | 0.96 | 0.98 |
|  |  |  | CHASE DB1 | 0.75/0.97 | 0.95 | 0.97 |
| Maji<sup>5</sup> | 2016 | CNN | DRIVE | - | 0.94 | - |
| Lahiri<sup>12</sup> | 2016 | CNN | DRIVE | - | 0.95 | 0.95 |
| Liskowski<sup>13</sup> | 2016 | CNN | DRIVE | 0.75/0.98 | 0.95 | 0.97 |
|  |  |  | STARE | 0.81/0.98 | 0.96 | 0.98 |
| Fu<sup>14</sup> | 2016 | CNN | DRIVE | 0.72/- | 0.94 | - |
|  |  |  | STARE | 0.71/- | 0.95 | - |
| Fu<sup>15</sup> | 2016 | CNN+ CRF layer | DRIVE | 0.76/- | 0.95 | - |
|  |  |  | STARE | 0.74/- | 0.95 | - |
|  |  |  | CHASE DB1 | 0.71/- | 0.94 | - |
| Melinscak<sup>16</sup> | 2015 | CNN | DRIVE | 0.72/0.97 | 0.94 | 0.97 |

### Retinal vessel detection from different database

Three types of datasets were used in the included studies. **Figure 7** evaluates the retinal vessel images detection images.

**figure 7:**
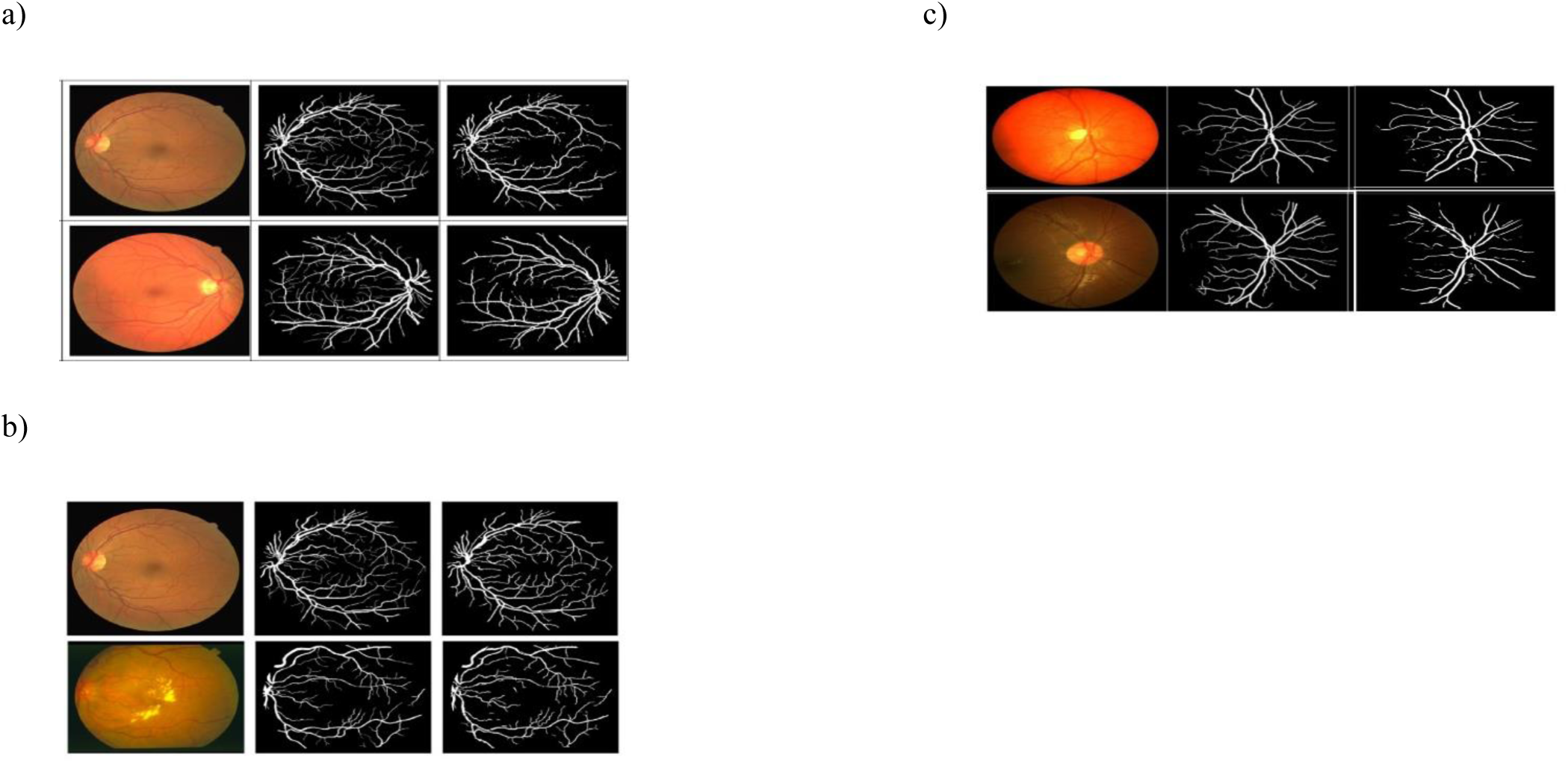
Visualization of the retinal vessel detection results from a) DRIVE dataset b) STARE dataset c) CHASE_DB1 dataset for original, corresponding and detection images.

### Conclusion

Although, the convolutional neural network algorithms are in their initial stages for detection of retinal vessel detection, but they have been shown some enthusiastic findings. The future role of CNN algorithms in the detection of the retinal vessel may be promising for its high accuracy and robustness. Automated detection of retinal vessel would help physicians or medical staff who are not very expert in retinal vessel detection imaging. It will contribute to making correct diagnosis and treatment faster and earlier. Integration of deep learning algorithms in real-world clinical setting would expectedly change traditional clinical practice for screening retinal images. With the increased prevalence of the retinal disease patients, deep learning based automated screening tools could ease the pressure on the healthcare system, particularly in under developing countries with large number of patients with eye diseases to be screen for eye disease with limited resources. Automated detection of retinal vessel may make the whole screening process more efficient, cost-effective and most importantly accessible to all.

